# Folate Receptor α Contributes to Radiation Resistance in Neuroendocrine Prostate Cancer by Regulating Redox Homeostasis

**DOI:** 10.64898/2026.03.26.714502

**Authors:** Hira Lal Goel, Tao Wang, Boris S Dimitrov, Ayush Kumar, Christi A Silva, Thomas J FitzGerald, Arthur M. Mercurio

## Abstract

Ionizing radiation can be an effective therapy for prostate cancer. Unfortunately, however, more aggressive prostate cancers such as neuroendocrine prostate cancer (NEPC) are often radiation resistant, which contributes to their high degree of morbidity and mortality. In this study, we used an unbiased approach to identify novel mechanisms that contribute to resistance to radiation and that are associated with neuroendocrine differentiation. Specifically, we compared the expression of cell surface proteins by mass spectrometry in prostate cancer cell lines that had been either untreated or treated with radiation to induce resistance, a process that also promotes neuroendocrine differentiation. Among the proteins identified by this screen, we focused on folate receptor α (FRα) because of its known biological functions and the fact that it is a validated therapeutic target. Our data reveal that FRα has a causal role in enabling prostate cancer cells to resist radiation. Importantly, we also demonstrate that the expression of FRα is regulated by HIF-1α, which also has a causal role in radiation resistance and neuroendocrine differentiation. Given that the ability of cells to resist damage and death in response to ionizing radiation depends largely on their ability to buffer the substantial increase in reactive oxygen species (ROS) that is generated by radiation, we also demonstrate that the folate-FRα axis promotes radiation resistance by sustaining intracellular glutathione levels that buffer this increase in ROS. In summary, the data reported here highlight a novel role for FRα in resistance to ionizing radiation that is intimately associated with the hypoxic microenvironment of NEPC and the ability of the folate-FRa axis to maintain redox homeostasis.

## Introduction

Advanced prostate cancer with metastasis is a disease that consists of multiple subtypes with significant heterogeneity that is primarily treated with androgen deprivation therapy (ADT). Although ADT is effective initially, it inevitably leads to resistance and the development of a lethal form of the disease termed metastatic castration-resistant prostate cancer (mCRPC) (Beltran and Demichelis, 2021). A subset of mCRPC can further develop into neuroendocrine prostate cancer (NEPC) by upregulation of regulators that drive stemness and lineage plasticity in response to pressure from therapies (Beltran and Demichelis, 2021; Li et al., 2025). NEPC most commonly evolves from mCRPC, and it is characterized by a stem-cell phenotype, expression of neuroendocrine markers, and loss or attenuation of androgen receptor signaling(Liu et al., 2022). It is an aggressive form of prostate cancer that is associated with rapid progression to metastasis, therapy resistance and a poor prognosis (Li et al., 2026). Currently, there are no effective therapies for treating NEPC. For this reason, there is urgent need to continue to understand the biology that underlies NEPC, and to exploit this knowledge to develop new treatment strategies to improve outcomes for patients.

Although ionizing radiation is a frequent treatment modality for prostate cancer, NEPC is highly radiation resistant, which contributes to its high degree of morbidity and mortality (Hu et al., 2015). Previous studies have investigated mechanisms that contribute to radiation resistance in NEPC (Hu et al., 2015). To date, multiple mechanisms have been described for how NEPC resists radiation therapy. These mechanisms include the observation that NEPC cells exhibit more efficient repair of radiation-induced DNA double-strand breaks and the observation that radiation selects for the survival of cells that grow more slowly and, consequently, are less susceptible to radiation-induced damage (Flores-Morales et al., 2019; Smith et al., 2015). There is also evidence that the hypoxic environment characteristic of advanced prostate cancer provides a survival advantage to more aggressive cells enabling them to resist radiation (Beckers et al., 2024). Despite the significance of these studies, a defined therapeutic target that sensitizes NEPC to radiation therapy has not been identified.

In this study, we sought to screen for specific cell surface proteins expressed on NEPC cells that contribute to radiation resistance, which could serve as potential therapeutic targets. To facilitate our approach, we took advantage of the observation that ionizing radiation can induce neuroendocrine differentiation in prostate cancer cell lines and patients. An unbiased mass spectrometry screen that compared the expression of cell surface proteins in prostate cancer cell lines that had been either untreated or treated with radiation to induce neuroendocrine differentiation revealed several proteins of interest whose expression is induced by radiation and associated with NEPC. Among these proteins, folate receptor α (FRα) caught our attention for several reasons. FRα is a GPI-anchored membrane protein that binds and transports folate, which is essential for DNA synthesis and cell proliferation, into cells in concert with other surface proteins including the reduced folate carrier and the proton-coupled folate transporter (Nawaz and Kipreos, 2022). It is overexpressed in many solid tumors such as ovarian cancer and, for this reason, FDA-approved therapies have been developed that inhibit its function (Moore et al., 2023). Importantly, a unique aspect of our study is the elucidation of a causal role for the FRα in radiation resistance. We also report that the mechanism by which the expression of FRα is induced is intimately associated with the hypoxic environment characteristic of NEPC.

## Results

### Radiation Induces Expression of FRα and Neuroendocrine Differentiation

We generated LNCaP cells that are more resistant to radiation than their parental counterparts by giving a total of 24 Gy over the course of several weeks using 4Gy x 6 doses treatment schedule (**Fig 1A**). These radiation resistant (RR) cells had an increased survival fraction following further exposure to radiation than parental LNCaP cells (**Fig. 1B**). Next, we verified previous reports that radiation induces neuroendocrine differentiation by assessing synaptophysin levels and observed that synaptophysin mRNA (**Fig. 1C**) and protein (**Fig. 1D**) were significantly higher in RR cells compared to parental cells. We also generated a radiation resistant PC3 model by treating PC3 cells with 8Gy of radiation x 6 doses treatment schedule (**Fig. 1E-F**) and observed that they too exhibited increased expression of synaptophysin mRNA (**Fig. 1G**) and protein (**Fig. 1H**).

**Figure 1.**
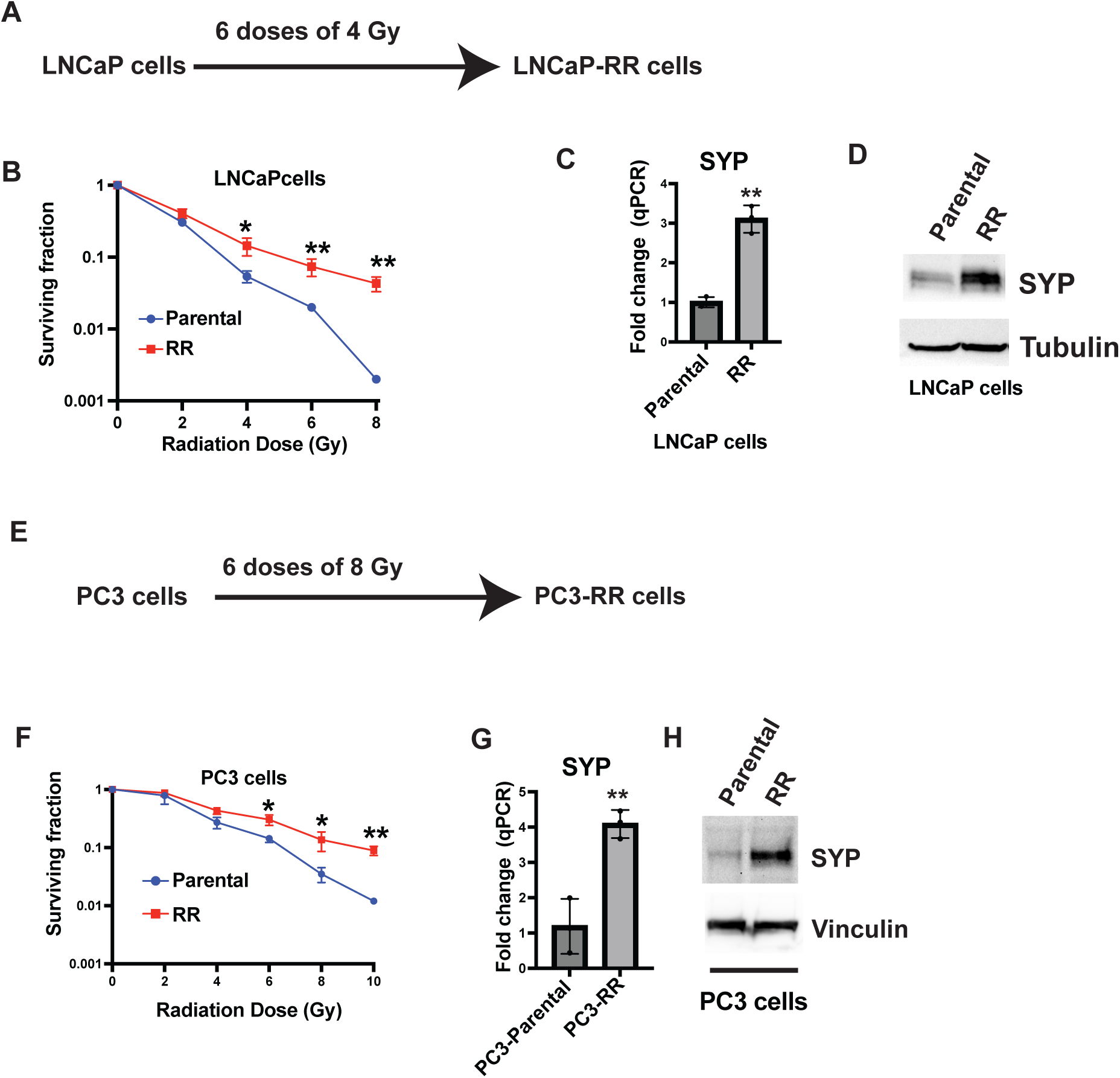
Generation of radioresistant cells with increased synaptophysin expression. **A.** Schematic representation showing generation of radioresistant LNCaP cells (LNCaP-RR). **B.** Clonogenic assay of LNCaP and LNCaP-RR cells comparing response to ionizing radiation (0-8 Gy). **C**. Synaptophysin mRNA levels were quantified in LNCaP and LNCaP-RR cells using qPCR. **D**. Synaptophysin protein was assessed in LNCaP and LNCaP-RR cells by immunoblotting. **E**. A schematic representation showing generation of radioresistant PC3 cells (PC3-RR). **F.** Clonogenic assay of PC3 and PC3-RR cells comparing response to ionizing radiation (0-10 Gy). **G**. Synaptophysin mRNA was quantified in PC3 and PC3-RR cells using qPCR. **H**. Synaptophysin protein was assessed in PC3 and PC3-RR cells by immunoblotting.

Our next goal was to identify specific cell surface proteins whose expression is induced in RR cells. To achieve this goal, surface proteins on parental and RR LNCaP cells were biotinylated and captured by immobilized avidin before being characterized by mass spectrometry. This approach revealed several surface proteins whose expression is induced by more than two-fold in RR cells (**Fig. 2A and Suppl Fig S1A**). Indeed, there were several strong candidates for subsequent analysis, including Plexin D1 and SLC22A3, a cation transporter. We focused our efforts on FOLR1 (FRα), however, for several reasons. Most importantly, antibody drug conjugates that target FRα are FDA-approved for ovarian cancer, supporting its feasibility as a potential therapeutic target in NEPC (Moore et al., 2023). Additionally, we observed increased mRNA expression of FOLR1 **(Fig. 2B**) in NEPC compared to non-NEPC specimens [available at cBioportal (Multi-Institute, Nat Med, 2016)]. As expected, increased expression of synaptophysin mRNA in NEPC specimens was also observed in the same set of patient samples (**Suppl Fig S1B**). We verified our mass spectrometry data by demonstrating that the expression of FOLR1 mRNA (**Fig. 2C-D**) and surface expression of FRα (**Fig. 2E**) are significantly higher in RR LNCaP cells and RR PC3 compared to parental cells. It has also been reported that FRα expression characterizes circulating tumor cells (CTCs) from prostate cancer patients (Lian et al., 2021). Furthermore, PSMA (prostate specific membrane antigen), an established biomarker for prostate cancer, helps to regulate folate absorption (Jajja et al., 2025). Despite these compelling indications, little is known about the function of FRα in NEPC or its potential contribution to radiation resistance.

**Figure 2.**
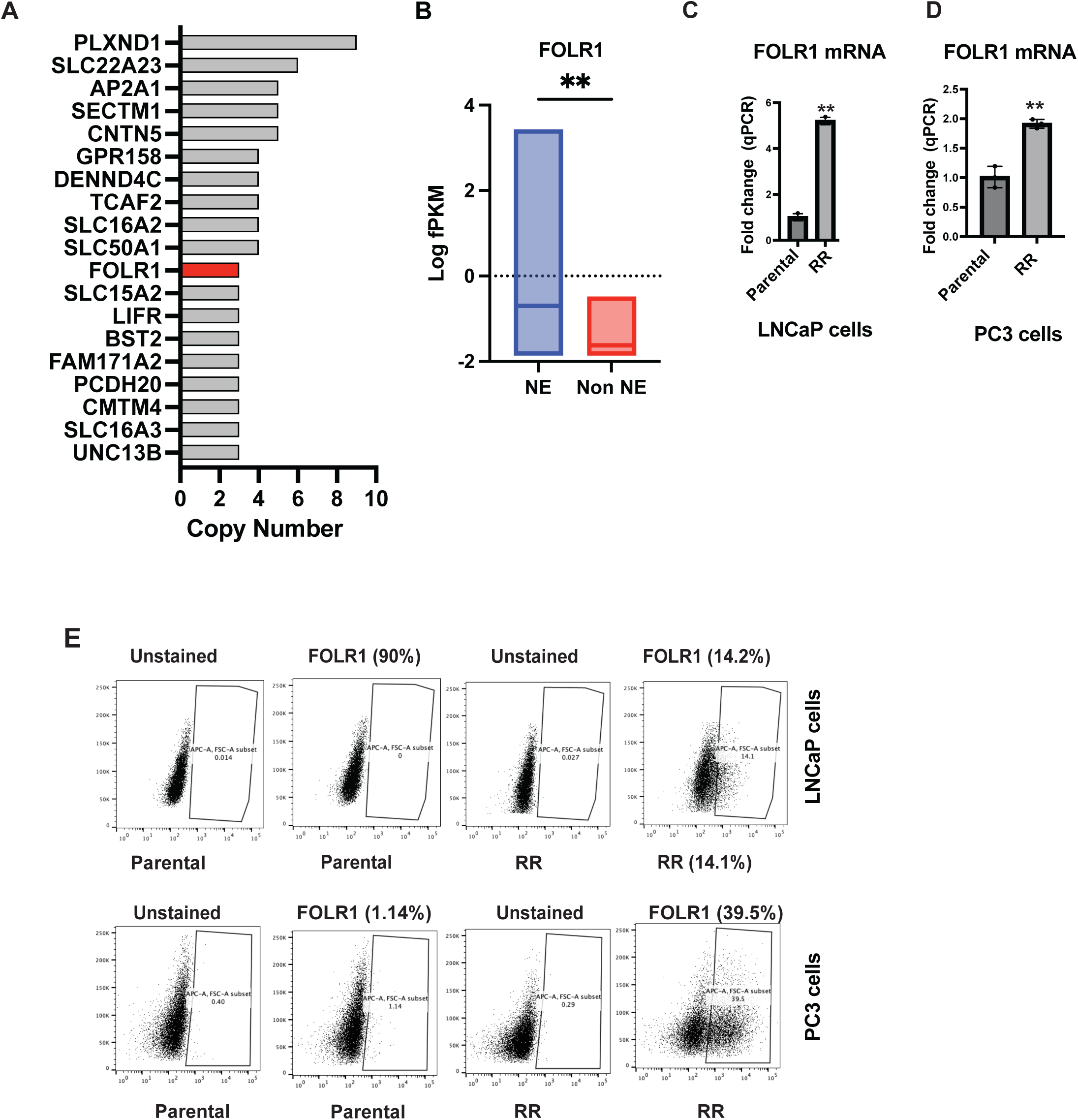
Resistance to radiotherapy is associated with increased expression of FOLR1. **A.** Total spectrum counts of differentially expressed surface proteins between control and radioresistant LNCaP cells is shown. **B.** FOLR1 expression counts (Log(Fragments per Kilobase million) were plotted using RNA-seq data from prostate tumors available at cBioportal. **C**. FOLR1 mRNA levels were quantified in LNCaP and LNCaP-RR cells using qPCR. **D**. FOLR1 mRNA levels were quantified in PC3 and PC3-RR cells using qPCR. **E**. Surface expression of FOLR1 was quantified in LNCaP, LNCaP-RR, PC3 and PC3-RR cells using flow cytometry.

### HIF-1a regulates FOLR1 expression

To investigate how radiation regulates the expression of FRα, we focused our attention on HIF-1α because of its previously reported function in the development of NEPC (Guo et al., 2019; Qi et al., 2010). Furthermore, radiation stabilizes HIF-1α expression which, consequently, has a major role in promoting radiation resistance (Moeller et al., 2004; Zhang et al., 2021). To confirm these observations, we verified that HIF-1α protein but not mRNA expression was elevated in LNCaP RR cells compared to parental cells (**Fig. 3A,B**). Similar increases in HIF-1α protein and mRNA levels were seen in PC3 cells (**Fig. 3C-D**). We also observed that knockdown of HIF-1α in LNCaP RR and PC3-RR cells decreased synaptophysin protein expression (**Fig. 3E and S2C**), consistent with previous reports on the role of HIF1α role in the development of NEPC. Importantly, knockdown of HIF-1α in LNCaP RR cells decreased FRα mRNA and surface expression (**Figs. 3F-G and S2A-S2B**). Similarly, PC3 RR cells had decreased expression of FOLR1 following HIF1 α knockdown (**Fig. 3H**). Conversely, exposure of these cells to hypoxia, a condition known to induce HIF1α, using CoCl_2_ increased FOLR1 and synaptophysin mRNA expression (**Figs. 3I and S2D-E**). However, inhibition of the FRα signaling using methotrexate, which inhibits DHFR, a downstream effector of FRα, or downregulation of FRα using shRNA did not affect the expression of synaptophysin protein or mRNA in either LNCaP (**Fig. 3J-K**) or PC3 cells (**Fig. 3L**) suggesting that this receptor does not contribute to NEPC differentiation.

**Figure 3.**
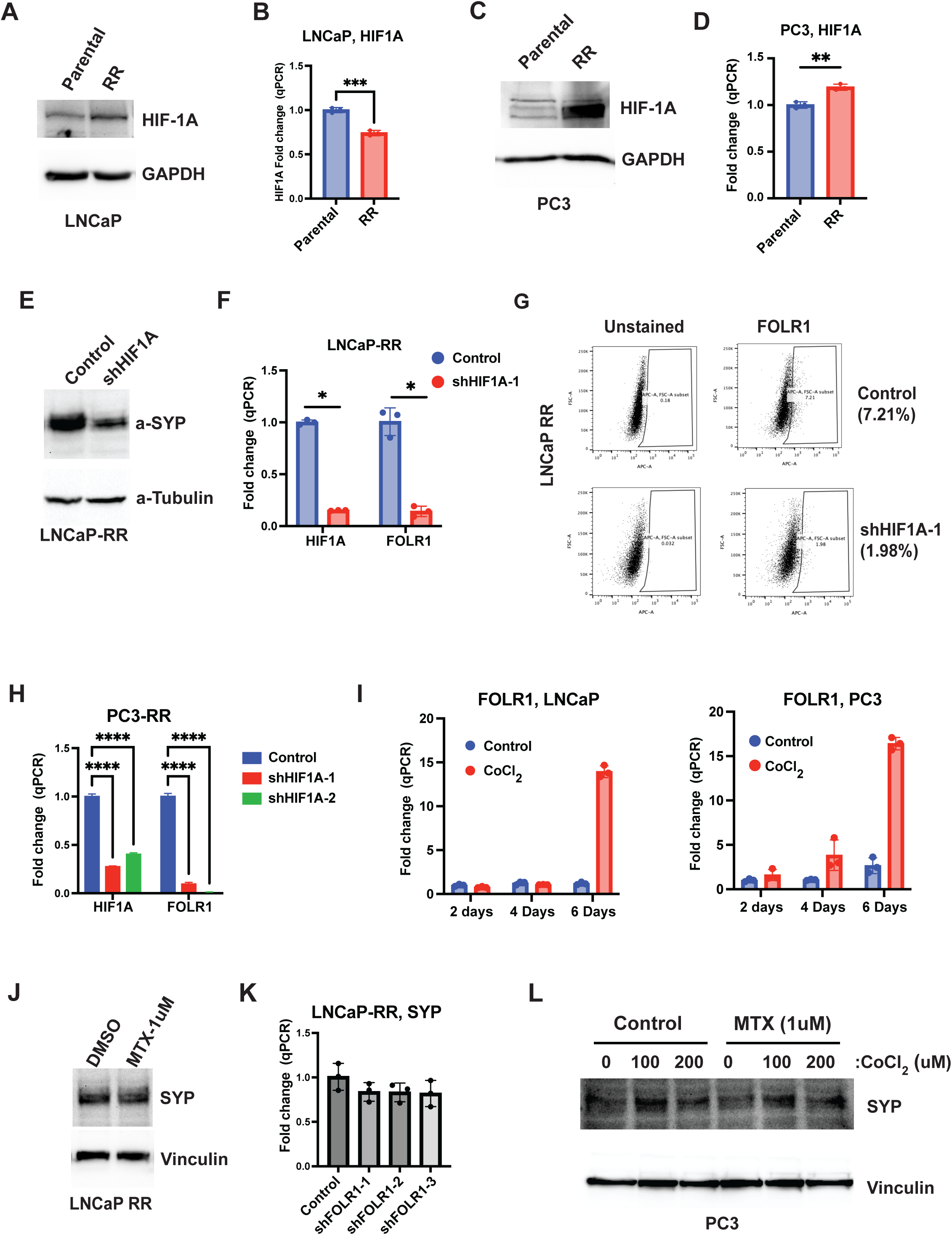
HIF-1α expression is essential for maintaining high FOLR1 expression in radioresistant prostate cancer cells. **A.** HIF-1α protein was quantified in LNCaP and LNCaP-RR cells using immunoblotting. **B**. HIF-1α mRNA was quantified in LNCaP and LNCaP-RR cells using qPCR. **C.** HIF-1α protein was quantified in PC3 and PC3-RR cells using immunoblotting. **D**. HIF-1α mRNA was quantified in PC3 and PC3-RR cells by qPCR. **E.** Synaptophysin protein expression was quantified by immunoblotting in LNCaP-RR cells after shRNA mediated downregulation of HIF-1α. **F.** HIF-1α and FOLR1 mRNA expression was quantified by qPCR in LNCaP-RR cells after shRNA mediated downregulation of HIF-1α. **G.** FOLR1 surface expression was quantified by flow cytometry in LNCaP-RR cells after shRNA mediated downregulation of HIF-1α. **H.** HIF-1α and FOLR1 mRNA expression was quantified by qPCR in PC3-RR cells after shRNA mediated downregulation of HIF-1α. **I.** FOLR1 mRNA expression was quantified by qPCR in LNCaP and PC3 cells after culturing cells in CoCl_2_ for 2, 4 and 6 days. **J.** Synaptophysin protein expression was quantified by immunoblotting in LNCaP-RR cells after culturing cells in 1 uM methotrexate for 24h. **K.** Synaptophysin mRNA expression was quantified by qPCR in LNCaP-RR cells after shRNA mediated downregulation of FOLR1. **L.** Synaptophysin protein expression was quantified by immunoblotting in PC3 cells after culturing cells in CoCl2 and methotrexate for 24 hours.

### FRα has a causal role in promoting radiation resistance

To assess a causal role for FRα in promoting radiation resistance, we either knocked down its expression in LNCaP RR cells using shRNAs (**Fig. 4A,B)** or treated these cells with methotrexate (**Fig. 4C**). Both approaches increased the sensitivity of these RR cells to increasing doses of radiation determined by a colony formation assay (**Fig. 4B, C**). Conversely, we expressed the FRα in parental PC3 cells exogenously, which resulted in a dramatic increase in its surface expression (**Fig. 4D**). This increased expression correlated with increased resistance to radiation (**Fig. 4E**). Lastly, we maintained both PC3 and LNCaP RR cells in folate-reduced medium for 72 hours to inhibit FRα signaling and then assayed their sensitivity to increasing doses of radiation. As shown in **Fig. 4 F,G**, cells maintained in folate-reduced culture medium were more sensitive to radiation than cells cultured in normal medium.

**Figure 4.**
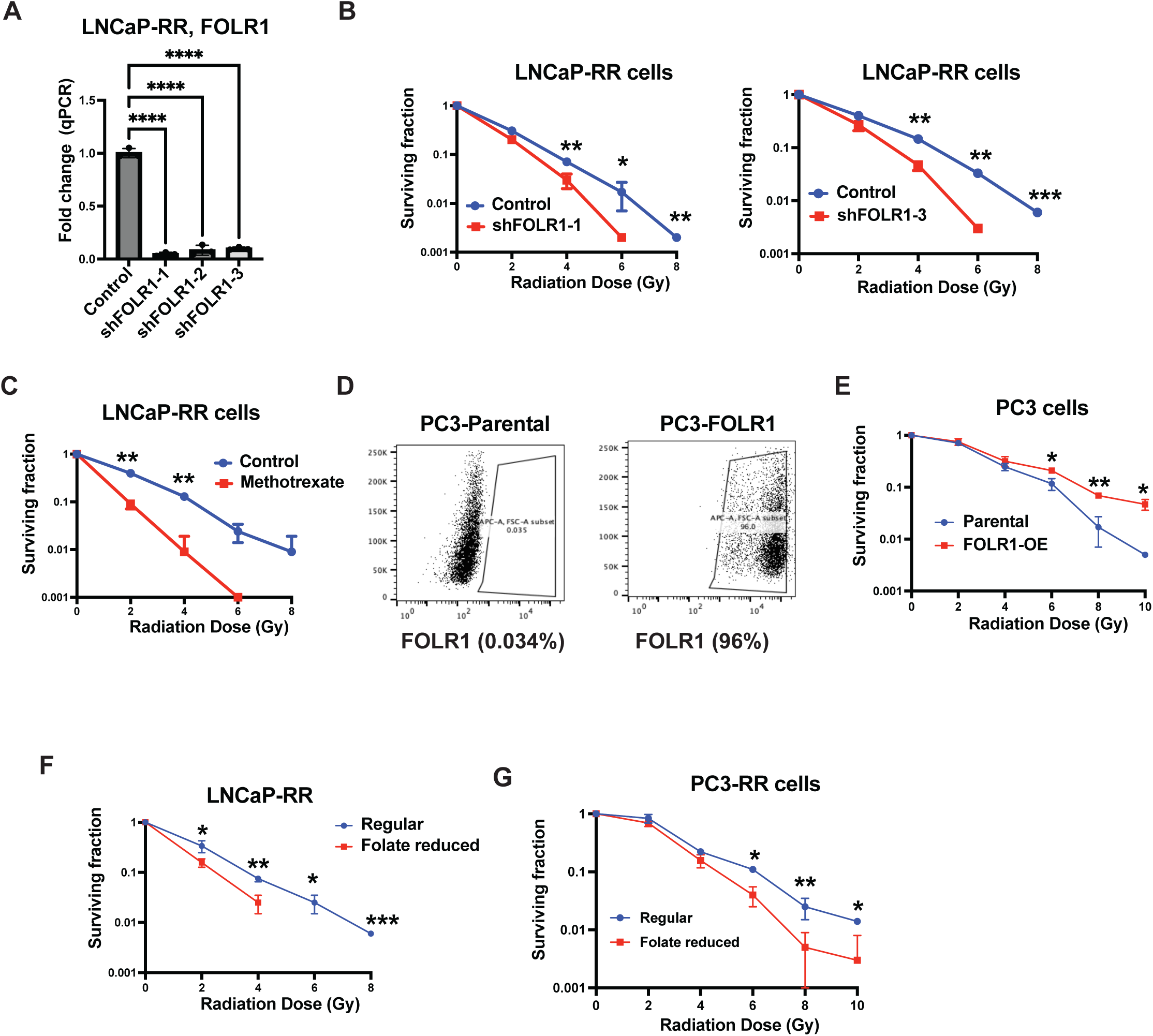
Folate/FRα signaling contributes to radiation resistance. **A.** FOLR1 mRNA expression was quantified by qPCR in LNCaP-RR cells after shRNA mediated downregulation of FOLR1. **B.** Clonogenic assay of LNCaP-RR control and shFOLR1 cells after irradiation (0-8 Gy) is shown. **C.** Clonogenic assay of LNCaP-RR cells treated with 1uM methotrexate after irradiation (0-8 Gy) is shown. **D**. Overexpression of FOLR1 in PC3 cells was confirmed by flow cytometry. **E.** Clonogenic assay of PC3 cells (control or overexpressing FOLR1) after irradiation (0-10 Gy) is shown. **F.** Clonogenic assay of LNCaP-RR cells performed under control or folate-reduced conditions after irradiation (0-8 Gy) is shown. **G.** Clonogenic assay of PC3-RR cells performed under control or folate-reduced conditions after irradiation (0-10 Gy) is shown.

### FRα promotes radiation resistance by enhancing GSH synthesis and ROS scavenging

Given our finding that the FRα has a causal role in promoting radiation resistance (**Fig. 4**), we investigated the mechanism involved. Radiation exposure induces the formation of reactive oxygen species (Wang et al., 2019), prompting us to investigate the possibility that the FR α might be involved in the generation of antioxidant defenses. This model is further supported by the fact that folate metabolism generates cysteine, a precursor for glutathione (GSH) biosynthesis, which serves as a major antioxidant that contributes to radiation resistance by maintaining redox balance (Lee et al., 2024). In support of this hypothesis, we observed that the LNCaP RR cells had higher levels of GSH than the parental cells (**Fig. 5A**). Knocking down FRα in these cells led to a reduced level of GSH relative to RR LNCaP cells (**Fig. 5B**). Moreover, knocking down FRα did not affect the expression of either GPX4 (the primary GSH utilizer) or SLC7A11 (the cystine/glutamate antiporter) (**Fig. S2F**), suggesting that the folate receptor-dependent increase in GSH is not due to altered expression of GPX4 or SLC7A11 and that the shift in antioxidant capacity is likely driven by the increased availability of cysteine via the folate cycle.

**Figure 5.**
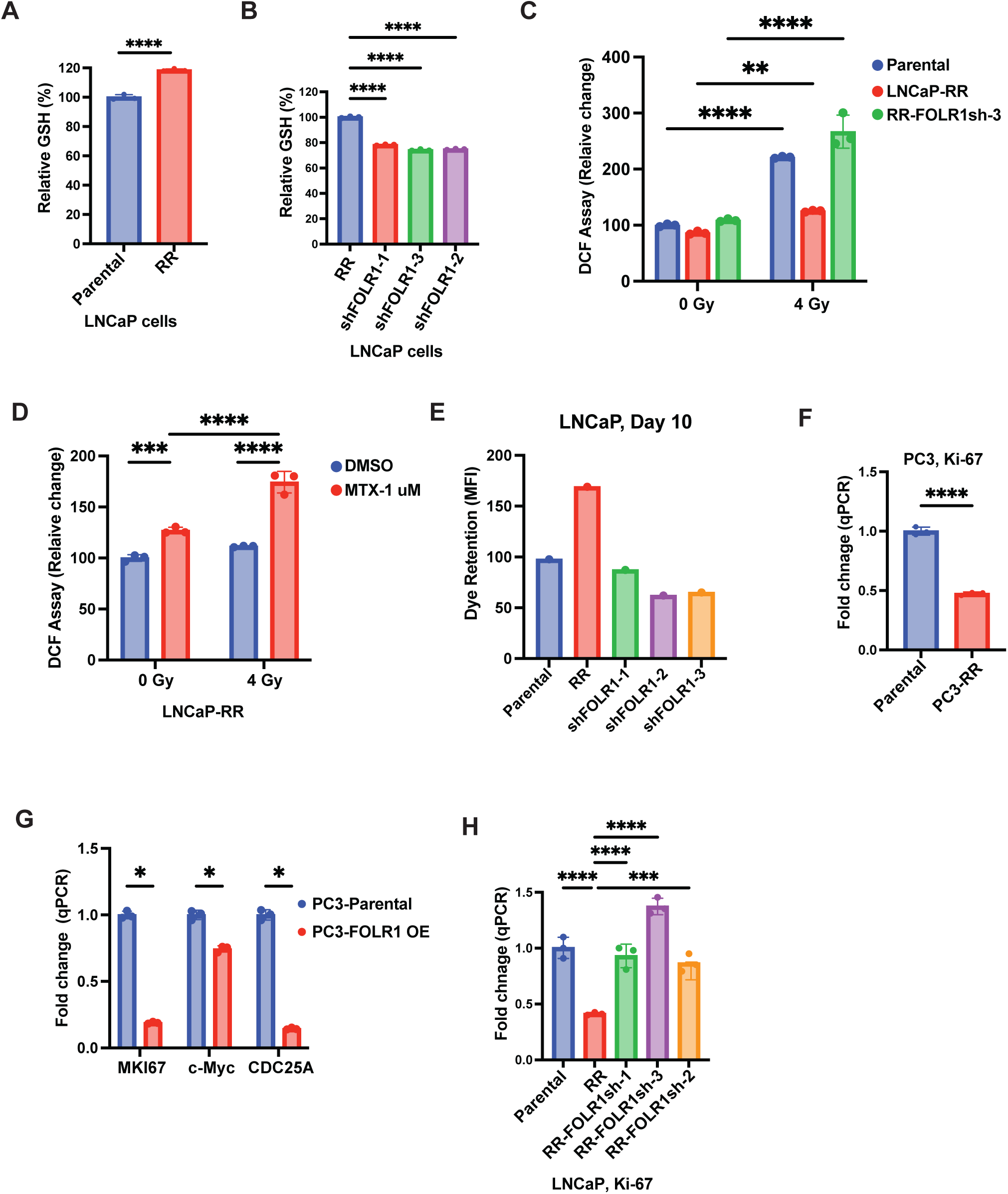
Folate/FRα signaling promotes radioresistance by enhancing GSH levels and reducing ROS. **A.** The levels of GSH were quantified in LNCaP and LNCaP-RR cells. **B.** The relative levels of GSH were quantified in LNCaP-RR cells after shRNA mediated downregulation of FOLR1. **C.** Relative levels of total ROS measured by DCF fluorescence is compared between LNCaP, LNCaP-RR and LNCaP-shFOLR1 at 24 hrs post radiation (4 Gy). **D.** Relative levels of total ROS measured by DCF fluorescence in LNCaP-RR cells cultured with or without Methotrexate (1uM) is shown 24 hrs after radiation (4 Gy). **E.** Bar graph plotted using mean fluorescence intensty to show retention of cell tracer violet dye in control, RR and RR with FOLR1 shRNAs populations of LNCaP cells. **F**. Level of Ki-67 mRNA levels was quantified by qPCR in control and PC3-RR cells. **G**. Effect of FOLR1 overexpression in PC3 cells on mRNA levels of Myc, Ki-67 and CDC25 was quantified by qPCR. **H.** Level of Ki-67 mRNA levels was quantified by qPCR in control, LNCaP-RR and LNCaP-RR-FOLR1shRNAs cells

Subsequently, we investigated whether the FRα-dependent GSH increase translates to enhanced ROS scavenging upon radiation. Under basal conditions, no significant differences in ROS levels were detected between parental and RR cells (**Fig. 5C**). However, radiation (4Gy) triggered an increase in ROS in parental cells but not in RR cells (**Fig. 5C**). To directly assess the contribution of the FRα directly to ROS regulation, we knocked down FOLR1 in RR cells and observed a significant accumulation of ROS following irradiation (**Fig. 5C**). In addition, treatment of RR cells with methotrexate mirrored the effects of FOLR1 knockdown, leading to a marked upregulation of radiation-induced ROS (**Fig. 5D**). We observed reduced cell growth in RR cells compared to non-irradiated cells (**Fig 5E**) and hypothesized the FOLR1 may be affecting cell growth and as a result diverting folate away from nucleotide synthesis towards GSH synthesis. To assess the effect of FRα on cell growth, we assessed the expression of cell growth related genes. We found that Ki-67 is significantly downregulated in PC3-RR cells and FRα expression in PC3 cells significantly inhibited the expression of Myc, Ki-67 and CDC25 (**Fig. 5F-G**). Similarly, Ki-67 expression is reduced in LNCaP-RR cells, which is rescued upon FOLR1 downregulation (**Fig. 5H**). Myc can sensitize cancer cells to oxidative stress because cysteine is utilized for protein synthesis instead of GSH generation and antioxidant defenses (Alborzinia et al., 2022; Dimitrov et al., 2025). Together, these results suggest that the folate-FRα axis contributes to radiation resistance by maintaining the GSH pool that buffers radiation-induced oxidative stress and slowing cell cycle progression.

## Discussion

The data reported in this study reveal a novel role for FRα in enabling prostate cancer cells to resist ionizing radiation, which is coupled with the known ability of radiation to induce neuroendocrine differentiation. We also report that HIF-1α, a key factor in the development of neuroendocrine differentiation that is associated with its hypoxic nature (Qi et al., 2010), regulates FRα expression. Importantly, we obtained data indicating that the ability of FRα to sustain GSH levels and, consequently, to buffer ROS is one mechanism by which it contributes to radiation resistance. Although FRα is associated with neuroendocrine differentiation, the data we obtained suggest that it does not have a causal role in this differentiation.

Although previous studies have reported that the expression of FRα is expressed on CTCs in prostate cancer patients, its functional and mechanistic contributions to prostate cancer have not been investigated extensively (Lian et al., 2021). Indeed, most studies on prostate and other cancers have focused on the role of folate because of its role in the one-carbon metabolism that is essential for DNA synthesis and proliferation. Interestingly, there are paradoxical effects of folate in prostate cancer depending on the context. For example, one prospective study reported that low folate intake was associated with an increased risk of recurrence in prostate cancer patients treated with radical prostatectomy (Tomaszewski et al., 2014) but another meta-analysis found that folate supplementation resulted in an increase in the relative risk of prostate cancer (Figueiredo et al., 2009; Wang et al., 2014). Also, increased one carbon metabolism has been shown to promote neuroendocrine prostate cancer (Reina-Campos et al., 2019). These studies, however, did not consider the contribution of FRα or other folate transporters. In this direction, several surface proteins, including the reduced folate carrier and the proton-coupled folate transporter can transport folate into cells in addition to FRα (O’Connor et al., 2021). We emphasize that our identification of the FRα as being associated with radiation resistance was based on an unbiased mass spectrometry screen of surface proteins whose expression is amplified in cells that are radiation resistant. Other folate transporters were not identified in this screen, indicating the specific role of the FRα in conferring radiation resistance.

We are particularly excited by our data implicating HIF-1α in the regulation of FRα expression because HIF-1α has a critical role in the development of NEPC, which is characterized by a hypoxic microenvironment (Qi et al., 2010). Moreover, our results suggest that HIF-1α regulates the transcription of FRα directly. Although other transcription factors including HNF4-α, Sp1 and AR, have been shown to regulate FRα transcription, the potential contribution of HIF-1α to its transcriptional regulation had not been considered (Salbaum et al., 2009; Sivakumaran et al., 2010; Yang et al., 2021). More than likely, HIF-1α functions in concert with other factors to regulate FRα transcription. Nonetheless, our data reveal that hypoxia and, presumably, hypoxic microenvironments in tumors, can increase FRα and its consequent functional effects.

Our findings also bear on the mechanism by which FRα contributes to radiation resistance. The ability of cells to resist damage and death in response to ionizing radiation depends largely on their ability to buffer the substantial increase in ROS that is generated by radiation (Wang et al., 2019). In this context, we were able to demonstrate that the folate-FRα axis sustains intracellular GSH levels and, consequently, buffers ROS. Although folate uptake is traditionally associated with rapid cell growth (Lee et al., 2024), our data reveal an unexpected and novel correlation between FRα upregulation and reduced proliferative capacity in radioresistant prostate cancer cells. Our finding suggests that under the selective pressure of ionizing radiation, FRα-mediated folate acquisition can be decoupled from the nucleotide biosynthetic machinery. Instead, these cells appear to employ a metabolic survival strategy wherein folate-derived one-carbon units are diverted toward the trans-sulfuration pathway to bolster GSH synthesis (Lee et al., 2024). This diversion to neutralize radiation-induced ROS comes at the functional cost of cell cycle progression as evidenced by reduced Ki67 and c-MYC expression in FRα-positive cells. Taken together, our findings suggest that cells can prioritize redox homeostasis over proliferation to survive radiation-induced ROS production and that the FRα can function not only as a growth promoter, but also as a metabolic rheostat that facilitates radio-resistance by reallocating resources toward the antioxidant defense system.

In summary, the data reported here highlight a novel role for FRα in resistance to ionizing radiation that is intimately associated with the hypoxic microenvironment of NEPC and the ability of the folate-FRα axis to maintain redox homeostasis. Given that FDA-approved FRα antibody drug conjugates are available, the possibility that these reagents or other therapeutic approaches to inhibit the FRα could be used to improve the response of patients with NEPC to radiation therapy merits consideration.

## Materials and Methods

### Cell Lines and Radiation

LNCaP cells (ATCC) were cultured in RPMI (Gibco, 22400-089) containing 5% FBS (Hyclone, SH3039603-SP), Sodium pyruvate (Gibco, 11360-070), MEM Non-essential amino acids (Gibco, 11140-050) and Pen/Strep (Gibco, 15140-122). PC3 cells were cultured in RPMI containing 10% FBS and Pen/Strep. To create folate-reduced conditions, cells were cultured for 3 days in folic-acid free RPMI with reduced serum (2% FBS) (Wu et al., 2023). Cells were irradiated using a 6 MeV Varian 2300CD linear accelerator (Varian Medical Systems, Palo Alto, CA). For colony formation assay we gave single doses of 2-10 Gy at 3 Gy/minute. Hypoxic conditions were induced by incubating cells with media containing CoCl2. We developed radioresistant models of LNCaP and PC3 by giving a total of 24 Gy (LNCaP) and 48 Gy (PC3) over the course of several weeks. Before moving on to the next radiation dose, we waited for the cells to reach 70 to 80% confluency in the plate.

### Reagents and Antibodies

We used following reagents for this study: Methotrexate (TargetMol, T1485), Folate-free RPMI (Gibco, 27016-021), CoCl2 (alfa Aesar, A16346). We used the following antibodies: Synaptophysin (Thermofisher, SP11, MA5-16402), GAPDH (Cell Signaling, 14C10, 2118S), HIF1alpha (Cell Signaling, 3716S), Vinculin (Abcam, Ab91459), FOLR1-APC (Miltenyi, 130-129-530) and Tubulin (TU-02, SC-8035).

### Plasmids and qPCR primers

The FOLR1 overexpression plasmid (pcDNA3.1-FOLR1) was purchased from Genscript (OHu14848D). The HIF1A shRNAs (pGIPZ; V3LHS_374854 and V2LHS_236718) and FOLR1 shRNAs (pLKO, TRCN0000060343, TRCN0000060344 and TRCN0000060346) were procured from shRNA Core facility at UMass. The primers for qPCR were obtained from Genewiz and sequences are:

**FOLR1** FW- GCTCAGCGGATGACAACACA

R-CCTGGCCCATGCAATCCTT

**SYP** FW- CTCGGCTTTGTGAAGGTGCT

REV- CTGAGGTCACTCTCGGTCTTG

**GAPDH** FW- GGAGCGAGATCCCTCCAAAAT

REV- GGCTGTTGTCATACTTCTCATGG

**HIF1A** FW- GAACGTCGAAAAGAAAGTCTCG

REV- CCTTATCAAGATGCGAACTCACA

**c-Myc** FW- GGCTCCTGGCAAAAGGTCA

REV- CTGCGTAGTTGTGCTGATGT

**CDC25a** FW: TCTGGACAGCTCCTCTCGTCAT

REV- ACTTCCAGGTGGAGACTCCTCT

**Ki-67** FW-ACGCCTGGTTACTATCAAAAGG

REV-CAGACCCATTTACTTGTGTTGGA

### DCF Assay

1×10^5^ cells per sample were detached, washed with PBS and incubated in 10uM 2’,7’-dichlorodihydrofluorescein diacetate (DCF) for 30 minutes in the dark at room temperature. Following incubation, cells were washed with PBS twice and measured for fluorescence (Ex/Em = 485/535nm) using a GloMax plate reader (Promega).

### Cell Proliferation Assay

Cell proliferation was assessed using CellTrace Violet Proliferation Kit (Invitrogen, C34557). Cells were harvested, washed twice with pre-warmed PBS and resuspended in 1 ml PBS. The staining dye was added to a final concentration of 5 uM and incubated for 20 minutes at 37C. To stop the staining, cells were incubated with 5 ml of complete culture media for 5 minutes. Cells were washed with PBS twice and resuspended in the complete culture media. One set of the cells were used for FACS analysis after incubating for 10 minutes (0 time point). The second set of cells were cultured for 7 days and processed for FACS (7 day time point). All data sets were analyzed using Flowjo software.

### GSH Assay

For total GSH levels, 3×10^5^ cells per sample were detached, washed with cold PBS and lysed using mammalian cell lysis buffer (M-PER Mammalian protein extraction Reagent, cat 78501 ThermoFisher). The GSH/GSSG Ratio Detection Kit (Abcam) was used according to the manufacturer’s instructions to quantify GSH. Fluorescence (Ex/Em = 490/520nm) was measured using a GloMax plate reader (Promega) and total GSH levels were calculated based on results from a linear regression analysis of the GSH standard curve.

### Mass Spectrometry

We used cell surface protein biotinylation and isolation kit from Thermoscientific (A44390) to biotinylate the surface protein in control and radioresistant LNCaP cell clones. These clones were generated by exposing cells to ionizing radiation at 5 Gy/ week for 5 weeks. After total of 25 Gy irradiation, the cells were kept in culture till surviving cells grew as colonies. The colonies were harvested and cultured as individual clone. We followed the instruction provide by the User Guide from this kit. The proteins were eluted using the elution buffer plus DTT and submitted to Mass spectrometry Facility at UMass Chan Medical School. Briefly, the samples were digested using S-trap and dried in speedvac. After drying, the samples were reconstituted with 20 µL of 0.1% Formic acid in 5% acetonitrile, followed by centrifugation at 16000 rcf for 15 minutes. Subsequently, 18 µL of the sample was transferred to non-binding HPLC vials.1 µL of the samples were injected and analyzed on a TimsTOF Pro2 (Bruker) mass spectrometer coupled to a nanoElute LC system (Bruker). An in-house packed C18 column (250 mm x 75 μm id, 3 μm, 120 Å pore size) was utilized with a 30-minute gradient. The flow rate was set at 500 nL/min and a solvent composition of Solvent A: 0.1% FA in water and Solvent B: 0.1% FA in ACN. The DDA data was searched against a reviewed human proteome downloaded from UniProt. The search was made using Fragpipe/MSFragger specific (trypsin) workflow.

### Flow Cytometry (FACS)

To measure surface expression of FOLR1, cells were detached and washed using complete media. 200,000 cells were incubated with a-FOLR1-APC antibody (1:100) for 40 minutes on ice. The cells were washed three times with PBS and pellet was resuspended in PBS. The samples were analyzed using BD FACS Celesta and data was analyzed using Flowjo software.

### Colony formation assay

Cells were detached, counted, plated in 6-well plate and irradiated. The number of cells per well varied depending upon the radiation dose. The cells were fed with fresh media after 72 hours. After 10 (PC3 cells) to 16 days (LNCaP), cells were fixed using 1% paraformaldehyde (in PBS) for 30 minutes. Cells were washed with PBS and stained using 0.5% Crystal violet for 1 hour. Colonies bigger than 50 cells were counted. The plating efficiency and the surviving fraction was calculated as described before (Kumar et al., 2024).

### Statistical Analysis

Student’s t test was used to compare between two groups. All statistical tests were carried out using GraphPad Prism 10.0 with a significance level set at *P* less than 0.05 and the following symbols represent the associated P-value: \**P*<0.05, \*\**P* <0.01, *** *P*<0.001, and **** *P*<0.0001.

## Acknowledgments

We thank Drs. Emmet Karner and Mengdie Wang and other members of Mercurio lab for critical discussions and cBioportal data analysis. This work was supported by NIH Grants 1R01CA276863 (AMM), R50 CA2211780 (HLG), and F30 CA275327-01A1 (AK). We thank Yansong Geng at Department of Radiation Oncology for irradiating cells.

---

**Figure S1.**
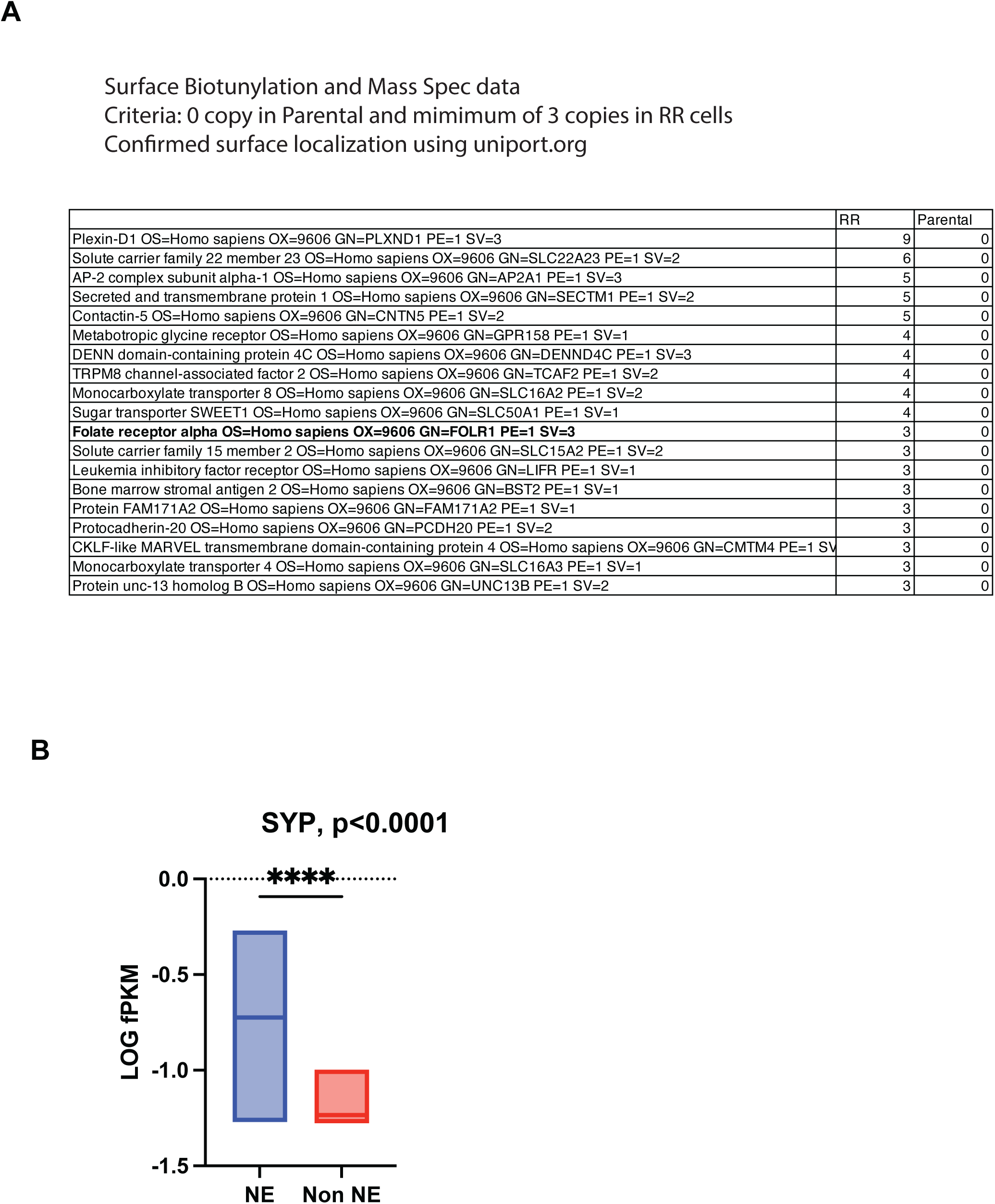
**A.** A list of differentially expressed surface proteins in LNCaP and LNCaP-RR cells with spectrum count is shown. **B.** Synaptophysin expression counts (Log fpkm) were plotted using RNA-seq data from prostate tumors available at cBioportal.

**Figure S2.**
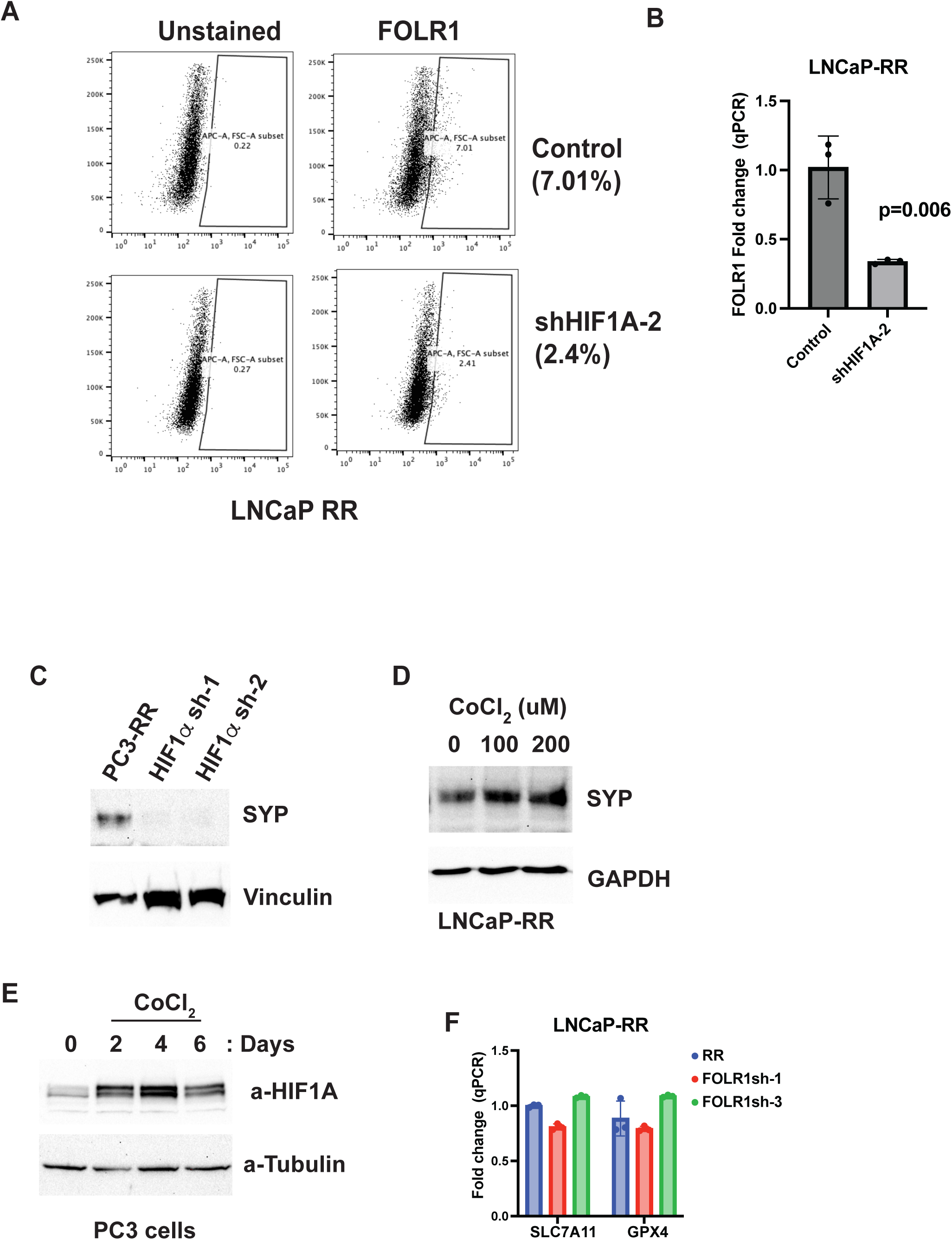
**A.** FOLR1 surface expression was quantified by flow cytometry in LNCaP-RR cells after shRNA mediated downregulation of HIF1A (shRNA-2). **B.** HIF1A mRNA expression was quantified by qPCR in LNCaP-RR cells after shRNA (shRNA-2) mediated downregulation of FOLR1. **C.** Synaptophysin protein expression was quantified by immunoblotting in PC3-RR cells after shRNA mediated downregulation of HIF-1α. **D.** Syp mRNA expression was quantified by qPCR in LNCaP-RR cells after culturing them in CoCl_2_ for 24 hours. **E.** HIF-1α protein levels were quantified by immunoblotting in PC3 cells after culturing cells in CoCl_2_ for 2, 4 and 6 days. **F.** SLC7A11 and GPX4 mRNA expression was quantified by qPCR in LNCaP-RR cells after shRNA mediated downregulation of FOLR1.

